# Prioritizing Neglected Food Species in Nutritional Studies Using Expert-Knowledge and Explainable AI

**DOI:** 10.1101/2025.03.26.645402

**Authors:** Michelle Cristine Medeiros Jacob, Aline Martins de Carvalho, Ângela Giovana Batista, Aníbal de Freitas Santos Júnior, Antonio Augusto Ferreira Carioca, Celia Márcia Medeiros de Morais, Cinthia Baú Betim Cazarin, Daniel Tregidgo, Danilo Vicente Batista de Oliveira, Dirce Maria Lobo Marchioni, Eliana Bistriche Giuntini, Elias Jacob de Menezes-Neto, Fillipe de Oliveira Pereira, Gabriela de Farias Moura, Hani R. El Bizri, Ingrid Wilza Leal Bezerra, Jailane de Souza Aquino, João Victor Mendes Silva, Josiane Steluti, Juliana Kelly da Silva-Maia, Juliana Araujo Teixeira, Lara Juliane Guedes da Silva, Letícia Zenóbia de Oliveira Campos, Marcela Alvares Oliveira, Maria Elieidy Gomes de Oliveira, Mariana de Paula Drewinski, Marina Maintinguer Norde, Nelson Menolli, Priscila F. M. Lopes, Rafael Ricardo Vasconcelos da Silva, Rômulo Romeu Nóbrega Alves, Samara Camile Gomes da Silva, Sávio Marcelino Gomes, Severina Carla Vieira Cunha Lima, Thais Q. Morcatty, Ulysses Paulino Albuquerque

## Abstract

Food biodiversity is vital for human health and sustainable food systems. However, research on neglected and underutilized species is limited by funding, uneven research capacity, and the challenge of balancing ecological, cultural, and public health considerations, requiring innovative prioritization approaches. Using Brazil as a model, this study inventories 369 neglected food species across algae, aquatic fauna, wild terrestrial vertebrates, insects, mushrooms, and plants. A mixed-methods approach, combining expert knowledge and explainable AI (LightGBM and SHAP value analysis), identified key factors for prioritizing species for food composition and consumption studies. The inventory is dominated by plants (29.5%) and wild vertebrates (24.4%), with major nutritional data gaps, particularly for algae, insects, and wild vertebrates. Over 36,000 recipes using neglected species were identified. In both food composition (R²: 0.677) and consumption studies (R²: 0.782) recipe number and species occurrence across different states were the most influential features predicting prioritization. These findings emphasize the role of cultural uses and local accessibility in shaping nutritional research priorities. We urge increased research on neglected species to bridge data gaps and integrate them into food systems, promoting sustainable diets in Brazil and other tropical regions.

## Introduction

Food biodiversity, defined as the diversity of plants, animals, and other organisms used for food, both cultivated and from the wild, plays a crucial role in global health and food security [1]. Within this context, neglected species, which include a range of wild and/or underutilized plants, fungi, algae, and wild animals (in specific contexts), represent an untapped resource for improving nutrition and sustainable food systems [2]. While no objective criteria definitively separate neglected from non-neglected food species, a general consensus defines neglected species as those whose contributions to sustainable food systems are significantly undervalued [3, 4]. This undervaluation stems from a widespread lack of awareness, limited information, and minimal presence in established markets.

The relevance of food biodiversity to human health and to food and nutrition security (FNS) is multifaceted. Biodiversity significantly influences food availability, quality, and stability — key components of FNS that directly impact human health outcomes [5]. In fact, increased Dietary Species Richness (DSR) improves micronutrient intake adequacy [6] and is associated with lower mortality rates, as shown by a large longitudinal cohort study conducted across nine countries [7]. Specifically, a 10-species increase in DSR was associated with a 14% to 17% reduction in all-cause mortality in men and a 6% to 8% reduction in women after an average 20-years follow-up.

These studies highlighted the association between higher dietary diversity and reduced mortality through four mechanisms [7]. The first, termed as “sampling effect”, posits that increasing DSR enhances the likelihood of including a wider array of highly nutritious and protective foods. The second, known as “complementary effect”, suggests that interactions between different food species can result in health benefits greater than the sum of their individual effects. The third mechanism, “minimizing trade-offs”, reduces the risk of toxicity from the overconsumption of any single species, even considering the natural presence of anti-nutritional factors or agrochemicals. Lastly, dietary diversity provides a broad range of nutrients and substrates, such as dietary fiber and bioactive compounds, that support the development of various gut microbes, fostering microbiome diversity. This change in gut microbiota, in turn, enhances health responsiveness, as a more diverse microbiome tends to be more stable and resilient to disturbances [8, 9].

In this context, food composition and consumption studies are crucial for shaping nutritional science and public health policies. Food composition studies systematically analyze and document food’s nutritional and chemical makeup, including macro- and micronutrients, bioactive compounds, and other essential components. The findings from these studies contribute to creating and maintaining food composition databases [10], which are fundamental for establishing dietary guidelines and informing nutrition policies. On the other hand, food consumption studies collect detailed data on the types and amounts of foods consumed by specific populations, tracking dietary habits, consumption patterns, and factors influencing food choices [11]. Therefore, food composition and consumption studies help identify the nutritional gaps and areas where dietary improvements are needed, making them indispensable tools in addressing malnutrition in all its forms and promoting public health. These two fields of study are deeply interconnected: food composition data is essential for accurately assessing nutrient intake, while food consumption data highlights which foods should be the focus of composition analyses [12].

Despite their importance, researchers face significant challenges in determining which neglected species should be prioritized for inclusion in food composition and consumption studies. Studies on food composition and consumption of neglected and underutilized food species face multifaceted challenges. The sheer diversity of species, uneven research capacity, and the complexity of integrating ecological, cultural, and nutritional priorities into a cohesive strategy complicate deciding what species should be the target of research efforts. Prioritizing species for research requires careful consideration of multiple factors, including nutritional value, cultural significance, ecological importance, and research needs, among others. These factors are exemplified in the literature and summarized in Box 1. These challenges highlight the urgent need for innovative, data-driven approaches to strategically prioritize species for study, ensuring that limited resources are effectively utilized to address critical gaps and unlock the potential of these species in advancing sustainable and healthy food systems.

#### Box 1. Criteria for Choosing Neglected Food Species in Nutritional Studies

##### Food Composition Studies

1. Research Gaps: Prioritize species with limited or outdated nutritional data [12].
2. Cultural Significance: Focus on culturally significant or traditionally consumed foods that lack comprehensive nutritional data [13].
3. Anecdotal or Preliminary Evidence: Consider species with anecdotal reports of health benefits or preliminary studies suggesting nutritional value that require further scientific examination [14].
4. Ecological and Agricultural Significance: Include species that play important roles in local ecosystems or agricultural systems but have yet to be thoroughly studied for their nutritional content [15].
5. Emerging or Underutilized Species: Prioritize species gaining popularity or those with potential for broader use that lack comprehensive nutritional information [4].
6. Biodiversity Conservation: Considers species at risk of genetic erosion or extinction to document their nutritional profile before potential loss while simultaneously implementing protection measures and sustainable management strategies to prevent overexploitation resulting from increased awareness and demand [16].
7. Climate Resilience: Include species known for adaptability to changing environmental conditions to understand their nutritional profile [17].

##### Food Consumption

1. Survey goal: Species that reflect the objective of the proposed dietary assessment, which can be tracking specific food groups or nutrients, a specific health outcome, or a whole dietary pattern. [3]
2. Local Availability and Accessibility: Focus on available species that are easily accessible to communities [18].
3. Cultural Significance: Prioritize culturally important or traditionally consumed species [13].
4. Underutilization: Include underutilized species to understand consumption patterns and the potential for promoting their intake. [2]

##### Overlapping Criteria

1. Cultural Significance: This section emphasizes the importance of studying foods with traditional or cultural values in a community. Including such foods in studies helps preserve cultural knowledge and potentially uncovers nutritional content.
2. Local Availability and Use: Prioritizes foods that are easily accessible to local communities. Studying locally available foods in both composition and consumption contexts ensures the research is relevant and applicable to the target population’s daily lives.
3. Underutilized or Emerging Species: This category focuses on foods that are not widely used but have the potential for greater utilization. By studying these in both composition and consumption contexts, researchers can identify nutritionally valuable foods that could contribute to dietary diversity and food security.
4. Research Goals and Gaps: This section emphasizes aligning the selection of foods with the overall objectives of the study, whether it’s filling knowledge gaps in nutritional data or understanding consumption patterns.

This article seeks to address the challenge of prioritizing neglected foods through two main objectives. First, we used Brazil as a model to compile and characterize an inventory of neglected food species in the country. Second, we utilized this inventory as a basis to develop and apply a data-driven framework for prioritizing food species for nutritional composition and consumption studies, integrating expert-knowledge and machine learning techniques. A thorough evaluation of the key factors driving species prioritization for nutritional composition and consumption studies can offer a replicable framework for guiding and informing effective resource allocation in both research and policy decision-making.

## Methods

### Study Area

Brazil is the largest tropical country in the world, and boasts one of the world’s wealthiest biodiversities, with an estimated 1.8 million species representing 9.5% of the global total [19]. However, this vast biodiversity wealth remains largely untapped and understudied, particularly regarding its potential contribution to food systems and nutrition. For example, a comprehensive study revealed that only 1% of Brazilians incorporate neglected and underutilized species into their diets, underscoring the enormous unexploited potential of Brazil’s food biodiversity [2]. Furthermore, research focusing on the Brazilian semi-arid, locally known as Caatinga, found that only 20% of its neglected plant species were represented in the country’s primary Food Composition Tables (FTC) [20]. Additionally, key species from various Brazilian biomes [21] are absent from the FCT most widely used by nutrition professionals in the country [22], such is the case of *baru* (*Dipteryx alata* Vogel) from the Cerrado, *carandá* (*Copernicia alba* Morong) from the Pantanal, *butiá* (*Butia odorata* (Barb.Rodr.) Noblick) from the Pampa, *araçá* (*Psidium cattleianum* Sabine) from the Atlantic Forest, and *camu-camu (Myrciaria dubia* (Kunth) McVaugh) from the Amazon, which is the richest known plant source of vitamin C globally [23]. Furthermore, no species of wild mushrooms, insects, or algae are represented in this FCT. These examples highlight not only the limited utilization of Brazil’s diverse food resources but also the significant gaps in our understanding of their nutritional composition, emphasizing the need for more comprehensive research and promotion of these underutilized species in Brazilian diets and food systems [24].

### Creation of Lists of Neglected Foods

A panel of experts was assembled based on their expertise in nutrition and environmental sciences and experience with the following food categories: algae, terrestrial wild vertebrate animals, fish and seafood, insects, mushrooms, and plants. Figure 1 outlines the entire research process, detailing the roles of all actors and steps involved.

**Figure 1.**
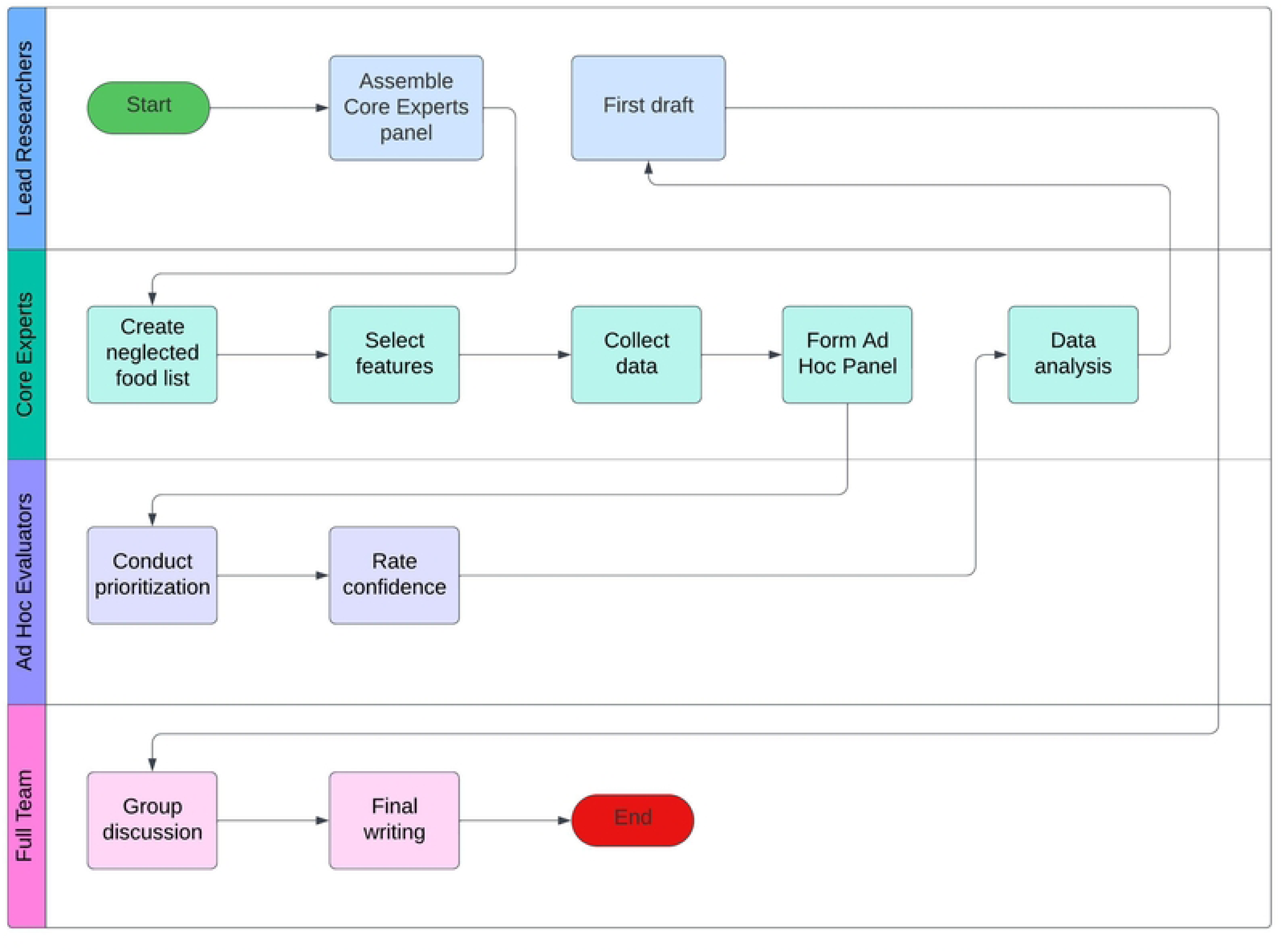
Study Workflow and Expert Involvement

These experts, referred to as ’core experts,’ were selected opportunistically from the lead researchers’ professional network. The panel also included one expert in data science. This team has been integral to the research since its inception, and they collaboratively compiled comprehensive lists of neglected food species across the six categories. These datasets were compiled following specific criteria to each specific food group (detailed in Supplementary Material 1), including only species with confirmed food use.

### Selection of Features and Source of Information

The core experts, alongside the lead authors, contributed to the selection of features used to characterize the neglected species. Since no predefined set of features existed, six were identified through a non-systematic literature review (see Box 1) and group discussions. The selected features were: regional occurrence in Brazil (number of Brazilian states with confirmed occurrence), conservation status with nine threat categories (*The IUCN Red List of Threatened Species*), origin (wild or cultivated), commercial cultivation status (Yes/No), availability of nutritional data (No; Yes, at the species level; Yes, at the genus level; Yes, by common name), and the number of recipes incorporating the ingredient. We implemented a structured approach to collect recipe information for each neglected food species, from the main sources Household Budget Survey (POF/IBGE), cookbooks that focus on neglected species and traditional Brazilian cuisine, and online sources, we chose the three most visited Brazilian recipe websites. Detailed sources for each type of information are provided in Supplementary Material 2. Data collection to populate the database with these features occurred between April and October 2023.

### Prioritization and Confidence Rating Process

Our project progressed to a species prioritization and confidence rating phase, which involved a separate group of researchers, referred to as ’*ad hoc* evaluators.’ These evaluators were specialists in either Nutritional or Environmental Sciences and had experience working with at least one of the neglected species categories covered in this study. To streamline recruitment, we developed a standardized form, which was distributed through the professional networks of the research team. All qualified specialists who expressed interest were included in the study. In recognition of their contributions, the evaluators were offered the opportunity to write and review the paper and participate as co-authors in the resulting publication.

In total, 18 *ad hoc* experts participated, including Nutritional Sciences (72.2%) and Environmental Sciences (27.8%), with a balanced distribution of specializations reflecting diverse backgrounds and areas of expertise. In Nutritional Sciences, the experts are evenly split between food composition and food consumption (38.5.8% each), with 23% experts in both fields. Among the food composition experts, many had experience across multiple food categories, with a significant proportion (83%) having extensive experience in plant-based foods, alongside expertise in other categories such as fish, fungi, algae, and insects. In Environmental Sciences, 60.0% of the experts specialize in ethnobiology, while 40.0% combine expertise in both ethnobiology and ecology. These specialists contribute diverse knowledge of plants, fish, and terrestrial wild animals. All experts declared no conflicts of interest. In our analysis, the variable “Labeler from nutrition science” highlights the differences in prioritization between specialists from Nutritional and Environmental sciences.

Before beginning the evaluation, *ad hoc* experts participated in a 30-minute orientation session to familiarize themselves with the task. They were also given a detailed dictionary of variables, which included comprehensive descriptions of each variable and, for categorical variables, outlined all possible response options. Then, they were provided with a spreadsheet containing 369 species, each described by six characterization variables (see previous section), along with two prioritization scales and two confidence calibration variables. To prevent bias, the species were not identified by their taxonomy; evaluators were only informed of the general category (algae, terrestrial wild vertebrates, fish, insects, mushrooms, or plants), in order to ensure that prioritization was based solely on species attributes. Using a Likert 1–5 scale (from very low to very high), the evaluators prioritized the species based on two key factors: their importance for nutritional composition studies and their relevance to dietary consumption research. They also rated their confidence in each of their responses on the same 1–5 scale.

Ethical approval for this study was not required under the Brazilian National Council of Health Resolution n. 510/2016, article 1, item VII. This resolution exempts research from Ethics Board review when no personal identification data is collected or published and when the information gathered pertains solely to the participants’ expertise rather than personal matters, as is the case of this study.

### Data Analysis

We examined the traits of the species included in the inventory using descriptive statistics, including metrics such as mean, median, standard deviation, and interquartile range to gain an understanding of the patterns and variability within the data. These data provide insightful information about a variety of characteristics, including habitat types, geographic distributions, and physical characteristics of the species.

We then used a supervised machine learning regression model to look into how these parameters affected the order of prioritization for composition and consumption research, and forecast prioritization scores according to the traits of the species. This method allowed us to capture and analyze the complex relationships between multiple independent variables (the species characteristics) and the dependent variable (the prioritization scores).

Then, we performed 10-fold cross-validation [25] to make sure our model was reliable and broadly applicable. This method divides the dataset into ten equally-sized pieces, or “folds.” After that, the model is trained ten times, with the training set consisting of the remaining nine folds and the validation set consisting of a different fold each time. By reducing variation and bias in the model’s performance estimation, this procedure offers a more accurate estimation of the model’s potential performance on unseen data.

Additionally, we selected the Light Gradient Boosting Machine (LightGBM) algorithm [26] as our modeling framework for the regression analysis. Using a gradient boosting architecture, LightGBM creates decision trees one after the other with the goal of fixing the mistakes of the previous ones. It is broadly used for its effectiveness and speed. We’ve selected LightGBM after assessing several commonly available architectures and detecting that LightGBM was the one with the lower Root Mean Squared Error (RMSE) in our data.

After training our final LightGBM model, we employed SHAP (SHapley Additive exPlanations) [27] to study the outputs from our model and comprehend how each feature affected the predictions. Employing ideas from cooperative game theory, SHAP values calculate each feature’s contribution to provide an explanation for a particular forecast. SHAP increases the transparency of the model’s decision-making process by giving each feature an importance value for a specific prediction.

To visualize SHAP data, we employed a variety of plots, such as force, dependence, and summary plots. With the use of these visualizations, we were able to determine not only which characteristics had the greatest overall influence but also how modifications to a certain feature’s value could impact the prioritization score. This degree of interpretability was essential to both the validation of our model and the provision of useful information for upcoming research.

We used several specialized Python libraries in our analysis. To process the datasets, we used Pandas [28]. We performed cross-validation, feature extraction and preprocessing using scikit-learn [29]. The gradient boosting algorithm came from the LightGBM package [30]. Using the SHAP library [31], we computed and displayed SHAP values. We assessed the model using traditional regression metrics, mainly the Root Mean Squared Error (RMSE), and R-squared (R²). These metrics offered numerical assessments of the model’s precision and capacity to account for the variation in the priority scores. The database and analysis scripts are available at this link https://github.com/eliasjacob/paper_bionut/

## Results

### Inventory of Neglected Food Species

We cataloged 369 species, with plants (29.5%) and wild animals (24.4%) predominating (see Supplementary Material 3). Mushrooms represented 23.6%, followed by fish and seafood (13.3%), insects (6.8%), and algae (2.4%). Almost half of the species (46.9%) lacked information on their conservation status, either because they had not been evaluated (44.4%) or were classified as data deficient (2.4%). Among those assessed, 42.8% are classified within the category least concern, while a total of 10.3% of the species face some degree of threat, being classified as critically endangered (0.3%), endangered (1.1%), vulnerable (5.7%), or near threatened (3.3%). The majority of species (81.6%) are native to Brazil. As for the commercialization, only 28.9% of the species are commercially cultivated.

Regarding food composition, just 33.1% of the species had available nutritional information (Supplementary Material 4). The group of foods that most frequently had nutritional information were plants, with 88.0% of the total species listed having available nutritional data. This is significantly higher than the nutritional information availability observed for other food groups, such as fish and seafood (24.5%), wild meat (15.6%), mushrooms (12.6%), insects (0%) and algae (0%). For the plant group, the nutritional information is predominantly available at the species level, with 62.4% of foods having species-specific data.

Out of the 36,058 recipes sourced in total (from the Household Budget Survey (POF) by the Brazilian Institute of Geography and Statistics (IBGE), cookbooks, and websites), only 72 preparations reported in the POF included ingredients from our list. The remaining recipes were sourced from cookbooks and culinary websites. Recipe distribution by food group was as follows: plants (60.3%), fish and seafood (20.5%), mushrooms (18.6%), and wild meat (0.6%). Recipes for algae and insects were scarce, with 10 recipes for algae and two for insects. *Manihot esculenta* Crantz (cassava) emerged as the species with the highest number of associated recipes—6,875, or nearly one-fifth of the total culinary references in the entire database (Supplementary Material 5).

### Factors that Predict the Prioritization in Nutritional Studies

In both food composition (R²: 0.677) and consumption studies (R²: 0.782) recipe number and species occurrence across different states were the most influential features predicting prioritization (Figure 2).

**Figure 2.**
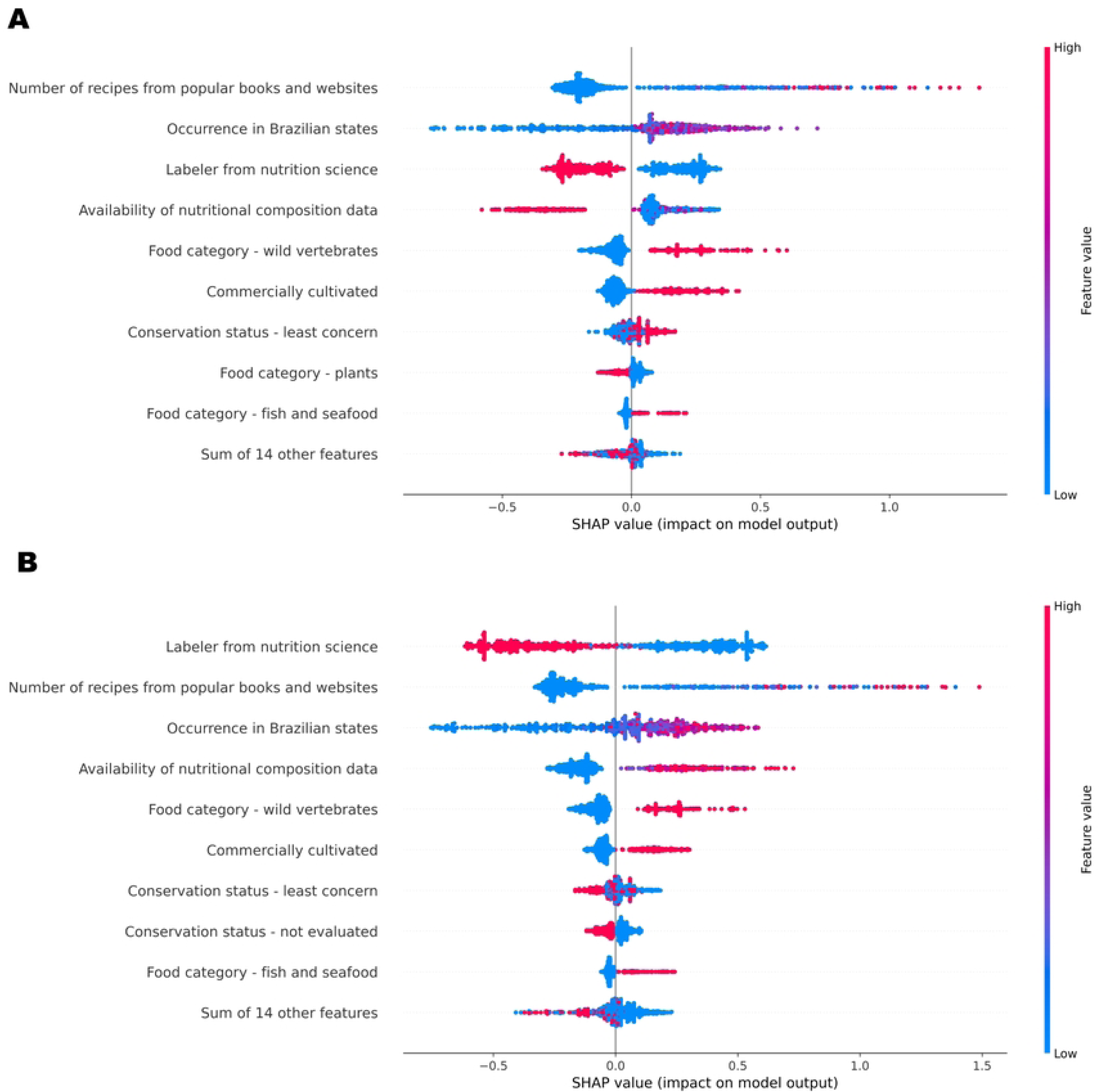
Priority factors for including species in studies of (A) food composition and (B) food consumption. The beeswarm plot uses SHAP (SHapley Additive exPlanations) values, where positive SHAP values (right) indicate that the feature is contributing to higher prioritization, and negative values (left) favors lower prioritization. Each point represents an observation in the dataset. The color gradient indicates the value of the feature for each observation, with blue points representing lower feature values and red points representing higher feature values. The SHAP values show the impact of each feature on the predicted prioritization score by the model.

Regarding the variables least associated with prioritization (i.e., the other 14 features mentioned in Figure 2), both in the context of composition and consumption studies, those related to species conservation status, as well as food categories such as insects and algae, exhibited the lowest influence. The variable “Labeler from nutrition science” highlights the differences in prioritization between specialists from nutritional and environmental sciences (Figure 2). Differences in feature-specific prioritization between specialist groups are summarized in Figure 3.

**Figure 3.**
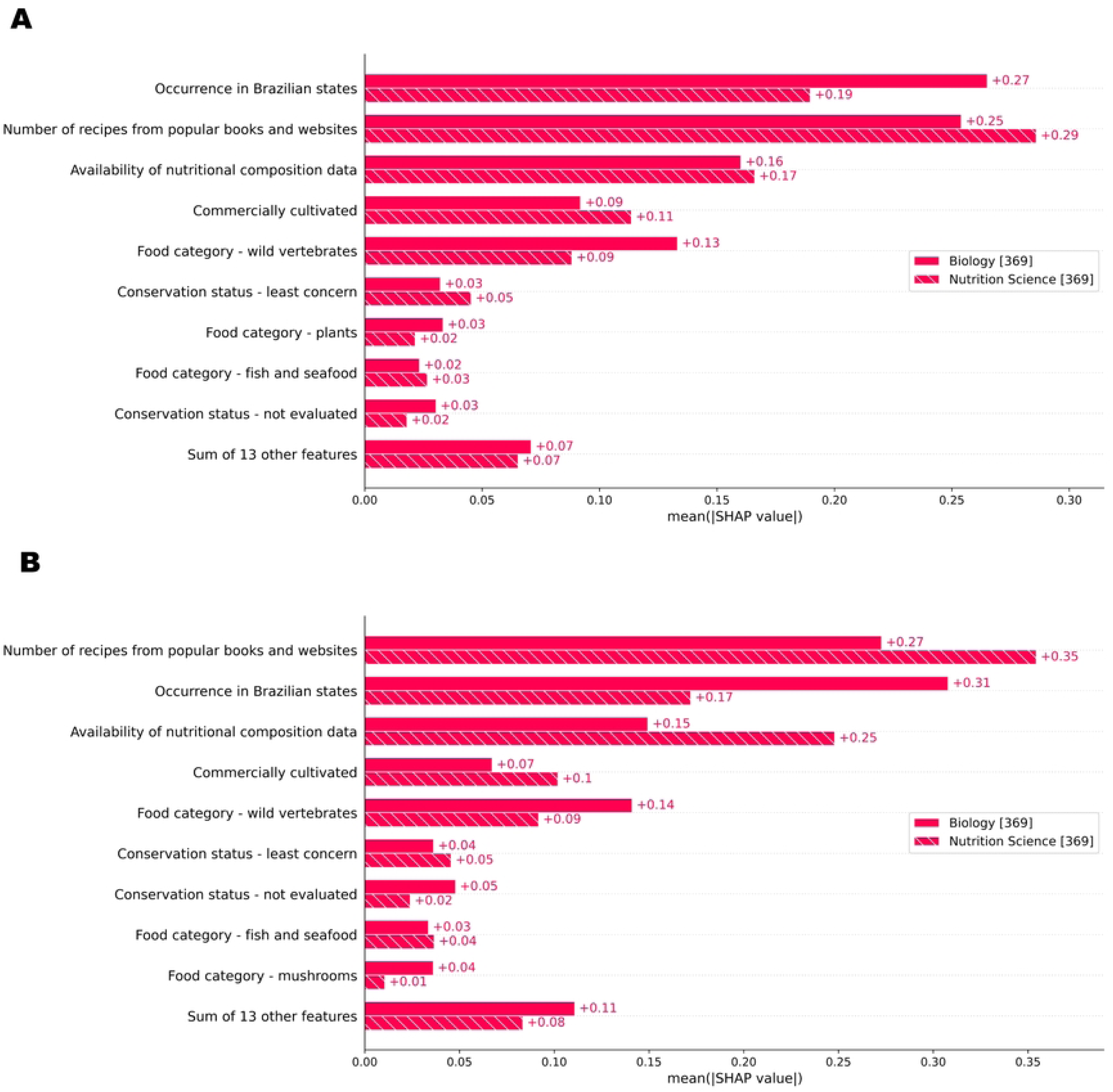
Comparison of Feature Prioritization Between Nutritional and Environmental Specialists.

Both specialist groups agreed on three key variables that should drive decision-making: a high number of recipes and a wide geographical distribution across Brazilian states (Figure 3). While there is a consensus that recipe number and species occurrence are the most important variables for prioritization, specialists differ in how they prioritize these factors. Environmental science specialists tend to prioritize species with a broader geographic distribution throughout the country, viewing this as the most important factor. In contrast, nutritionists place more emphasis on species with documented culinary uses, as evidenced by their presence in recipes.

While our study identifies key variables and their relative importance for prioritizing species, it doesn’t result in a simple mathematical formula for prioritization. This is because our analysis utilizes a LightGBM model, a complex machine learning algorithm that combines multiple decision trees to make predictions. Extracting a single formula from this intricate structure is impractical and wouldn’t accurately represent the model’s decision-making process. To ensure replicability, we provide access to our trained model and data on GitHub. Researchers can apply this model to their own datasets, using it to predict species prioritization based on the identified key variables. This approach offers a more nuanced and accurate assessment than a simplified formula could provide.

## Discussion

This study addresses the global challenge of prioritizing neglected food species by using Brazil as a case study to develop a broadly applicable framework. Our research identified 369 species, primarily plants and wild animals, many of which lack comprehensive nutritional data. The prioritization process revealed culinary applicability and distribution are the most influential criteria for selecting species. Ecological or conservation aspects were not key factors either, indicating a potential disconnect between nutritional research priorities and broader biodiversity conservation goals. Disciplinary perspectives also played a role, with environmental scientists prioritizing species with wider distribution. These findings underscore that species prioritization for composition and consumption studies is primarily driven by immediate practical utility and researchers’ expertise, potentially overlooking crucial ecological considerations. These insights underscore that nutritional studies require a more integrated strategy that balances immediate research needs with long-term ecological sustainability.

The prevalence of plants in our inventory reflects their historical dietary importance to early populations in Brazil and reveal a rich and diverse history of plant use, underscoring their long-standing role in people’s diets. Archaeobotanical evidence from the Cerrado region indicates that indigenous people utilized a variety of wild plants and later incorporated domesticated species such as maize, manioc, and beans, reflecting a transition from hunter-gatherer to more settled agricultural practices [32]. This historical reliance on plants is further supported by a systematic review on the use of wild food plants by Brazilian communities, which found that early populations extensively used wild plants, with fruits, leaves, and seeds accounting for 85.8% of plant parts consumed [33]. The accessibility of plants likely contributes to their prominence in diets, as opposed to algae, insects, and wild animals. A nationwide study of Brazilian dietary habits revealed that unconventional food plants (UFPs) are consumed more frequently than edible mushrooms and wild meat. Notably, edible algae and insects were excluded from the analysis due to insufficient or non-existent consumption data, highlighting significant gaps in the utilization of these potential food sources [2]. This plant-centric focus, while highlighting diversity within the plant group, also reveals significant knowledge gaps in other food groups, particularly insects and algae, underscoring the need for more comprehensive research into these understudied food sources.

Building on this understanding of plant prevalence, our study estimates that developing recipes and increasing the availability of nutritional composition data can heighten scientists’ interest in neglected species from local biodiversity, thereby contributing to their popularization and scientific knowledge development. The promotion of edible species such as algae, mushrooms, and insects could enhance dietary diversity within primarily plant-based sustainable diets [5]. The benefits of integrating food biodiversity beyond plants into sustainable diets include increased resilience of the food system, better alignment of food cultures with environmental contexts, income generation for local communities, and the revival of cultural knowledge [34, 35]. However, despite their consumption in many traditional communities, cultural barriers - such as fear of poisoning, familiarity, and food neophobia - continue to limit the acceptance of these food groups by the broader population [36, 37].

A key example of cultural barriers to sustainable diets in Brazil is the consumption of insects, a practice that remains marginal despite the country harboring the highest diversity of edible insect species in South America. A recent study identified 141 edible insect species in Brazil, with 140 having documented records of consumption, though many of these records are anecdotal or date back to the colonial period [38]. In urban areas, insect consumption is typically driven by curiosity for the “exotic” or by enduring cultural traditions, such as the consumption of ants from the Atta genus, known locally as “tanajura” or “içá”. Some edible species, however, are not consumed in Brazil despite being eaten in other parts of the world, highlighting the potential for expanding this practice. However, significant cultural and regulatory challenges impede broader acceptance of edible insects in Brazil. Insects are often perceived as exotic and evoke neophobia and disgust among Brazilians, particularly among older women with less formal education, while younger, more educated men are somewhat more receptive [39, 40]. Additionally, the absence of specific regulations on insect consumption raises food safety concerns and complicates their integration into mainstream diets [41]. These cultural and institutional barriers present significant obstacles to leveraging insects as a sustainable food source, beyond their occasional use by Indigenous peoples and local communities.

Wild mushrooms, like insects, are largely overlooked in the Brazilian diet despite their nutritional and medicinal value. A recent study identified 409 species of wild edible mushrooms (WEM) in Brazil, but only 86 have robust records based on molecular data or Brazilian nomenclatural types, highlighting the need for further studies to confirm their identity and occurrence [42]. Of these species, 93 are consumed in Brazil, 52 have uncertain or incomplete evidence of consumption, and 264 are not consumed, presenting a potential new food resource. While interest in mushroom foraging is growing, proper training in identifying edible species is crucial to prevent misidentifications and potential poisoning.

Fish and seafood face unique challenges due to inconsistent identification and labeling practices. Many species are sold under generic names, such as “pescada,” which can refer to three distinct species in our inventory, obscuring biodiversity and sometimes resulting in the unintentional consumption of threatened species [43]. This ambiguity complicates assessments of consumption patterns and poses conservation concerns. Furthermore, the nutrient quality of fish and seafood depends heavily on accurate species identification [44]. Addressing these challenges will require precise labeling practices and comprehensive studies to promote sustainable use of aquatic food resources.

The reliance on recipe numbers as a primary indicator for species inclusion in nutritional studies demonstrates the need for a more nuanced understanding of cultural food practices, with recipes serving as a valuable source of information. In this regard, our study reveals a significant discrepancy in the documentation of neglected and underutilized foods between national dietary surveys and popular culinary resources. The Brazilian Household Budget Survey reports a mere 0.2% presence of these foods, whereas popular websites feature over 36,000 recipes. This stark contrast underscores the limitations of traditional dietary assessment methods, such as 24-hour recalls and food frequency questionnaires, which often underestimate the consumption of episodically consumed foods like wild, forest, and underutilized species [45]. This finding emphasizes the urgent need for more comprehensive methods to collect, analyze, and report dietary intake data that accurately reflect the contribution of neglected foods to diets. Given the complexity of developing national surveys focused on food biodiversity in a country as diverse as Brazil, a strategic approach could be to initially focus on communities that rely on traditional food systems, which are recognized for their rich food diversity [46]. This targeted approach would provide valuable insights into consumption patterns, allowing for the refinement of evaluation tools and the initiation of food composition analyses for these resources. By addressing these gaps, we can break the cycle in which these foods are overlooked due to assumptions about their lack of consumption and excluded from studies due to insufficient compositional data. Beginning with these communities could yield crucial insights and lay the foundation for broader national efforts to integrate food biodiversity into dietary assessments.

This study reveals a generally aligned approach to species prioritization for food research between nutritionists and environmental scientists, with both groups recognizing the importance of recipe prevalence and data availability. However, nuances exist in their perspectives. Nutritionists tend to prioritize species with more recipes available, while environmental scientists may consider broader geographic distribution as an indicator of ecological significance. Despite these complementary viewpoints, a concerning gap emerges in the lack of emphasis on conservation factors. This oversight may partly stem from the limited availability of conservation data, as our study found that almost than half of the species lacked information on their extinction risk. This data gap likely contributes to the low importance given to conservation in species prioritization. While practical considerations are important, neglecting the ecological and conservation status of food species risks undermining the long-term sustainability of food systems [47]. Studies should particularly emphasize the interconnected roles of biodiversity in supporting both nutritional health and ecological sustainability. For instance, biodiversity loss in fisheries in the Peruvian Amazon has reduced the availability of key micronutrients like zinc and iron, underscoring the importance of conserving diverse species for both dietary stability and ecosystem health [48]. Conversely, promoting consumption of species at risk can harm their populations and ecosystems if not properly managed. To ensure that future decisions about species prioritization are informed by a comprehensive understanding of their ecological impact, it is crucial to prioritize the collection and integration of conservation status data, such as extinction risk assessments, into food biodiversity research.

This study presents several strengths, including the creation of a comprehensive inventory of food biodiversity in Brazil and the innovative application of machine learning techniques (LGBM and SHAP Value) to analyze species prioritization factors. The inventory provides a valuable baseline for future research, while the analytical approach offers insights into the decision-making processes of experts across disciplines. However, it also has limitations. Primarily, the prioritization analysis, though informative, relies on expert opinions that may not fully encapsulate the complexity of food systems. Furthermore, while extensive, the inventory may not be exhaustive and could benefit from supplementation with specialized studies focusing on specific regions or food groups. For example, El Bizri et al. [49] assessed the impact of illegal sport hunting on Brazil’s wildlife, documenting the hunting of 157 native species, including 19 endangered ones. Their study revealed significant wildlife depletion, particularly in the Atlantic Forest and Caatinga biomes, with a shift from large mammals to smaller birds as hunting pressure increased. A nutrition study focused on wild meat could leverage our inventory by providing essential species data, particularly for underutilized and threatened species. Each food group, such as wild meat or seafood, may require additional data from specialized studies to fully capture its complexity. This supplementary information would enhance the overall value of our inventory for biodiversity and food consumption research.

We recommend several strategies to promote the consumption of underutilized foods, with practical and policy implications. First, encouraging investment in interdisciplinary and collaborative nutritional composition studies of local biodiversity foods is essential to increase their visibility and validate their nutritional value. Second, expanding dietary assessments among communities with limited market integration—who likely consume more diverse diets—could provide valuable insights into traditional dietary practices and support the preservation of food biodiversity. Third, fostering the development of an integrated national database on the culinary uses and nutritional composition of local foods, while strengthening existing initiatives, would be instrumental in making this knowledge more accessible to researchers, policymakers, and the general public. Finally, promoting the inclusion of biodiverse foods in public programs and training professionals working with human nutrition can further support their adoption, aligning dietary diversity with broader public health and environmental sustainability goals. These actions can help bridge the gap between biodiversity conservation and dietary practices, building a more resilient and culturally adaptive food system. Finally, integrating biodiverse foods into public programs and training nutrition professionals are crucial steps toward aligning dietary diversity with public health and environmental sustainability goals. However, it is essential to consider the conservation status of each species and implement measures to prevent ecological impacts. This includes promoting certified agroecological and agroforestry production and educating consumers on the importance of purchasing from sustainable sources.

## Conclusion

This study provides a crucial foundation for understanding and leveraging Brazil’s rich food biodiversity for improved nutrition and sustainable food systems. By compiling a comprehensive inventory of neglected food species and analyzing expert prioritization factors, we have highlighted key areas for future research and policy interventions. Our findings reveal a significant bias towards species with readily available culinary applications, often overlooking the ecological and cultural significance of underutilized food sources. This underscores the urgent need to shift from a purely nutritional perspective to a more holistic approach that integrates ecological, cultural, and social dimensions of food biodiversity. Prioritizing research on underrepresented food groups like algae, insects, and wild mushrooms, particularly those with limited conservation data, is crucial. Furthermore, integrating traditional knowledge from indigenous peoples and local communities, who possess invaluable insights into the sustainable use of these resources, is essential for developing culturally relevant and ecologically sound food systems. Ultimately, bridging the gap between biodiversity conservation and dietary practices requires a multi-faceted approach. This includes investing in interdisciplinary research, expanding dietary assessments to capture traditional foodways, developing comprehensive food databases, and promoting the inclusion of biodiverse foods in public programs. These efforts bridge the gap between biodiversity conservation and dietary practices, fostering a more resilient, culturally adaptive, and ecologically responsible food system.

## Disclosure statement

The authors report there are no competing interests to declare.

## Funding

This research was supported by the High-Performance Computing Center at UFRN (NPAD/UFRN) and the ‘Conselho Nacional de Desenvolvimento Científico e Tecnológico’ – CNPq through a research grant to MCMJ (402334/2021-3) and AMC (444588/2023-0). The authors also thank CNPq for the research productivity scholarships awarded to EJMN (302582/2023-1), MCMJ (306755/2021-1), NMJ (314236/2021-0), and PFML (302365/2022–2). PFML acknowledges the funding from the Romanian Ministry of Research, Innovation and Digitalization for project JUST4MPA (760054, within the framework of PNRR-III-C9-2022— I8). AMC, MPD, and NMJ acknowledge the support of the Sao Paulo Research Foundation – FAPESP by, respectively, the following grants: 2022/03091-6, 2017/25754-9, and 2018/15677-0. HREB was supported by USAID agreement with CIFOR-ICRAF. The study received a contribution from the INCT Ethnobiology, Bioprospecting, and Nature Conservation, certified by CNPq, with financial support from the Foundation for Support to Science and Technology of the State of Pernambuco as a grant [Grant number: APQ-0562-2.01/17] given to UPA.

## Contribution Statement

MCMJ led the project with responsibilities in Conceptualization, Data curation, Formal analysis, Funding acquisition, Methodology, Project administration, Validation, Visualization, and Writing (original draft and review & editing). UPA provided Conceptualization, Supervision, and Writing (review & editing). EJMN contributed to Conceptualization, Formal analysis, Methodology, Visualization, and Writing (review & editing). AMC, DT, DVBO, DMLM, FOP, GFM, JVMS, LJGS, MPD, NMJ, PFML, RRVS, SMG, SMGS contributed to Conceptualization and Writing (review & editing). AGB, AFSJ, AAFC, CMMM, CBBC, EBG, HREB, IWLB, JSA, JS, JAT, LZOC, MAO, MEGO, MMN, RRNA, SCVCL, TQM contributed to Validation and Writing (review & editing). All authors have reviewed and approved the final manuscript and agree to be accountable for all aspects of the work.

## Supplemental Material List

Supplemental Material 1 - Criteria for Creating the List of Neglected Foods

Supplemental Material 2 - Features and Source of Information

Supplemental Material 3 - Inventory of Neglected Species

Supplemental Material 4 - Data on Nutritional Composition of Neglected Species Available in Food Composition Tables in Brazil

Supplemental Material 5 - Recipes Containing Neglected Species

## Graphical Abstract

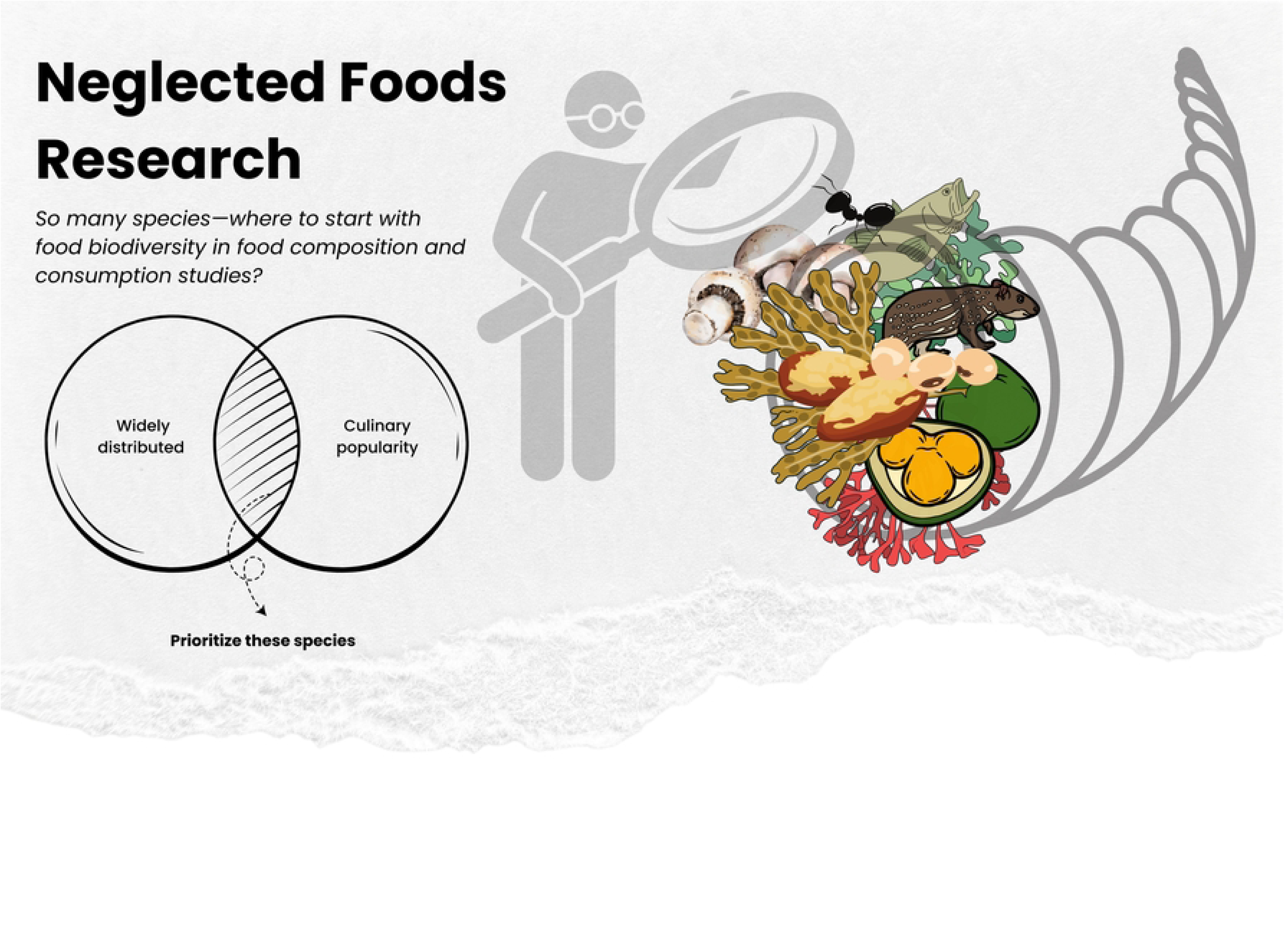

## Notes

### Competing Interest Statement

The authors have declared no competing interest.

